# Construction of simplified microbial consortia to degrade recalcitrant materials based on enrichment and dilution-to-extinction cultures

**DOI:** 10.1101/670133

**Authors:** Dingrong Kang, Samuel Jacquiod, Jakob Herschend, Shaodong Wei, Joseph Nesme, Søren J. Sørensen

## Abstract

The capacity of microbes degrading recalcitrant materials has been extensively explored from environmental remediation to industrial applications. Although significant achievements were obtained with single strains, focus is now going toward the use of microbial consortia because of advantages in terms of functional stability and efficiency. While consortia assembly attempts were made from several known single strains, another approach consists in obtaining consortia from complex environmental microbial communities in search for novel microbial species, genes and functions. However, assembling efficient microbial consortia from complex environmental communities is far from trivial due to large diversity and biotic interactions at play. Here we propose a strategy containing enrichment and dilution-to-extinction cultures to construct simplified microbial consortia (SMC) for keratinous waste management, from complex environmental communities. Gradual dilutions were performed from a keratinolytic microbial consortium, and dilution 10^−9^ was selected to construct a SMC library. Further compositional analysis and keratinolytic activity assays demonstrated that microbial consortia were successfully simplified, without impacting their biodegradation capabilities. These SMC possess promising potential for efficient keratinous valorization. More importantly, this reasoning and methodology could be transferred to other topics involving screening for simplified communities for biodegradation, thus considerably broadening its application scope.

**Importance:** Microbial consortia have got more and more attention and extensive applications due to their potential advantages. However, a high diversity of microbes is likely to hide uncontrollable risks in practice specific to novel strains and complicated interaction networks. Exploring a convenient and efficient way to construct simplified microbial consortia is able to broaden the applied scope of microbes. This study presents the approach based on enrichment and dilution-to-extinction cultures, which gain abundance microbial consortia including some without losing efficiency from the enriched functional microbial community. The microbial interactions at the strain level were evaluated by using compositional identification and correlation analysis, which contribute to revealing the roles of microbes in the degradation process of recalcitrant materials. Our findings provide a systematic scheme to achieve optimizing microbial consortia for biodegradation from an environmental sample, could be readily applied to a range of recalcitrant materials management from environmental remediation to industrial applications.

## Introduction

Microbial degradation aims to harness the potential of enzymatic processes of naturally occurring microorganisms to break down complex, and usually environmentally recalcitrant materials (1). Biodegradation is widely used in agricultural and industrial fields, especially when dealing with substances, which are difficult to dissolve and/or resistant to decomposition under mild conditions (2, 3). Large number of microorganisms have been isolated and characterized for their efficient capacity to degrade recalcitrant molecules such as organophosphorus compounds, being the most commercially favored group of pesticides, also posing a risk to human health (4, 5). Another example is the utilization of microbes to metabolize lignocellulose materials which represent a renewable carbon source with potential application as biofuel (6). To that end, microbial consortia gained from complex environmental communities harboring unknown genes, metabolic activities and species, hold a colossal application potential for enhancing the efficiency of bioprocesses, particularly when dealing with substances that are resistant to decomposition (7, 8). They have received increasing attention due to their promising advantages on handling environmental problems (9).

Microbial consortia consist of several species working together, providing both robustness and broad metabolic capacities (9). They may be assembled synthetically from scratch by combining several isolated strains (10), or conversely yielded from complex environmental microbial communities, like soil (11). Mixed populations are not only able to perform some difficult tasks that would not even be possible for individual strains, but offer more stability and resilience against environmental fluctuations (12). They are superior compared to single strains with respect to degradative efficiency in many cases (13). Unlike single strains isolation, enrichment process involving directed artificial selection is often used to obtain desired microbial consortia from the environment (14, 15). In general, composition of consortia gradually changes over time, including a richness reduction along with emergence of dominant microbes due to competitive exclusion (16). Nevertheless, many species are likely to benefit from the enrichment due to complex microbial interactions conferring functional stability and redundancy (17, 18). For instance, a total of 17 bacterial strains and 13 methanogens were identified from established microbial consortia depolymerizing lignin (19). Therefore, this intrinsic high diversity level of enriched microbial consortia represents a bottleneck in our attempts to move forward with potential industrial applications due to several aspects, like i) potential negative correlation with efficiency (20), ii) species with undesired function, iii) security threat posed by pathogens presence, and iv) risks of losing the properties of interest if hold by keystone species. Utilization of microbial consortia with less complexity, but equal efficiency, lead to more controlled and optimized industrial processes (10). Hence, it is crucial to find reliable strategies to narrow down the diversity towards optimum simplified microbial consortia. A reductive-screening approach was applied to construct effective minimal microbial consortia for lignocellulose degradation based on different metabolic functional groups (10). Additionally, artificial selection approaches (dilution, toxicity, and heat) have been employed to obtain desired bacterial consortia (21). A minimal effective hydrogen-producing microbial consortium was constructed *via* dilution-to-extinction culture from cow rumen communities (22). Dilution-to-extinction culture is expected to provide more advantages compared to conventional isolation and assembly as it i) generates many microbial combinations ready to be screened, ii) includes all strains from the initial microbial pool that might be lost due to cultivation/isolation biases and iii) ensure that all microbes are physically present and interacting spontaneously (23).

Keratins are insoluble fibrous proteins with cross-linked components, representing the most abundant proteins in epithelial cells (24). Keratinous materials are classified as one of the waste materials categories according to European parliament regulations (25). They could cause an environmental imbalance and potential pollution to soil or water because of their recalcitrant form in municipal waste (26). Hence, seeking the effective approach to keratinous waste management will contribute to eliminate their environmental risks. In fact, keratinous materials are estimated to have considerable economic value after biodegradation (27). A microbial consortium (KMCG6) was enriched from an environmental sample culturing in α-keratin based medium, which possesses an efficient keratinolytic activity (11). Despite successful diversity reduction during enrichment, it still includes plenty of microbes affiliating to distinct genera (11). In this work, we applied the concept of dilution-to-extinction culture to KMCG6 as a case study (Fig. 1), resulting in efficient simplified microbial consortia (SMC).

**Fig. 1.**
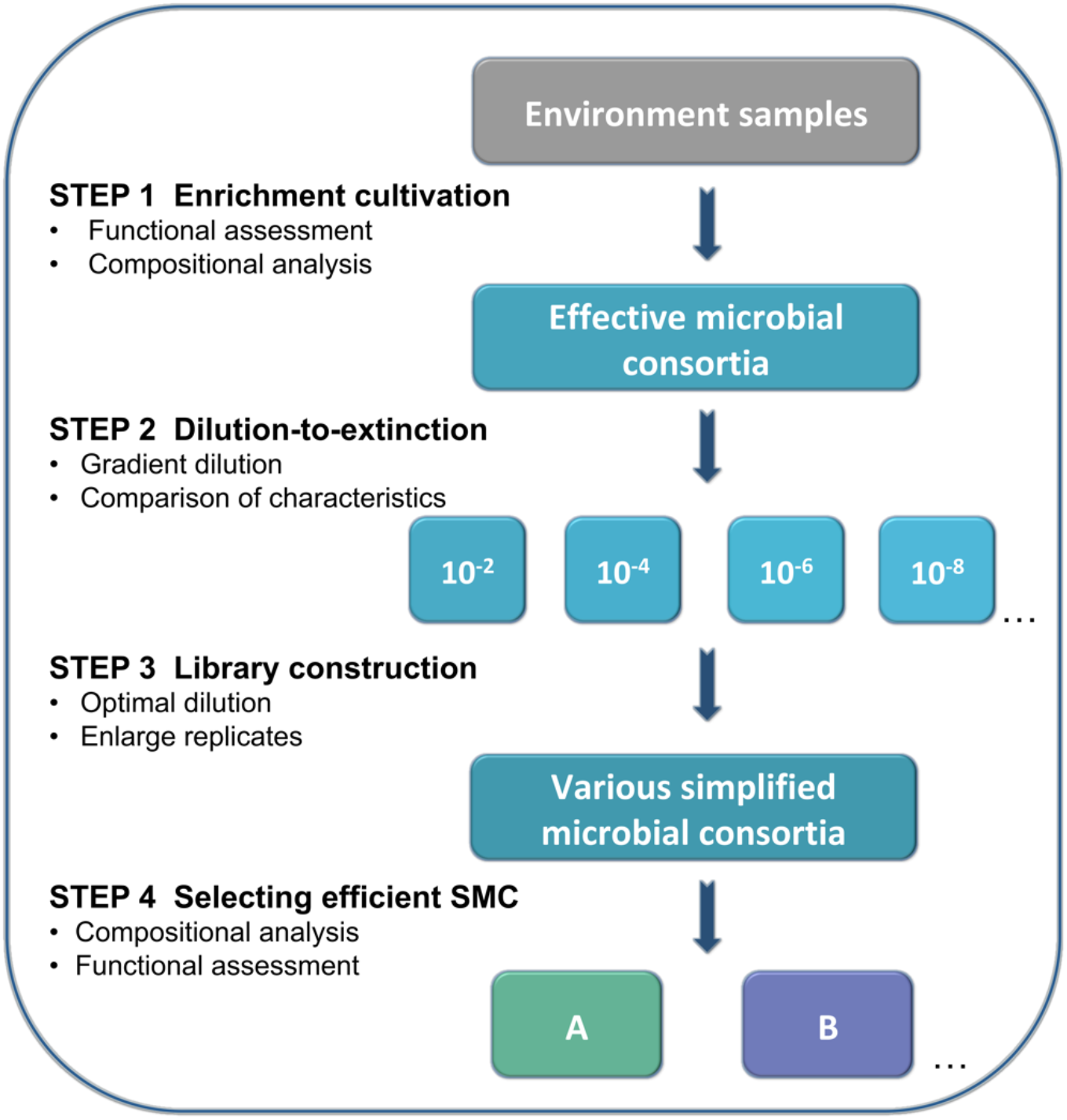
Workflow of enrichment and dilution-to-extinction cultures to select simplified microbial consortia (SMC) for keratin degradation. This workflow includes four steps: 1) Enrichment for the desired traits e.g. keratinolytic activity by selection in keratin medium, where keratin is the sole carbon source. Several serial transfers were performed to obtain effective microbial consortia. This process was evaluated by functional assessment (cell density, enzymes activity, and ratio of the residual substrate) and compositional analysis. 2) Gradient dilution was conducted to the enriched effective microbial consortia. Six gradual dilutions were prepared, from dilution 10^−2^ to 10^−10^ with 24 replicates. The dissimilarity between dilutions was evaluated by Euclidean distance calculation based on functional assessment criteria. 3) Library construction was done from the dilution offering the optimal dissimilarity among replicates. Dilution 10^−9^ was selected to construct the SMC library in this case. 4) Selection of the most promising SMC is based on functional and compositional characterization.

## Results and Discussion

### Phylogenetic analysis of microbial consortium KMCG6

Microbial consortia have shown numerous advantages for the processing of recalcitrant molecules and materials which are hard to degrade or convert (9), and may even represent environmental and threats (26). Microorganisms represent a colossal, yet still a poorly tapped resource for applied microbiology and biotechnology. Many efforts attempting to learn from nature have used microbial consortia for bioconversion. Microbial consortium KMCG6, displaying efficient keratinolytic activity, was enriched from a soil sample as previously described (11). Taxonomic composition analysis at phylogenetic levels (class, genus, and OTU) along the enrichment process was done over time at different generation batches (KMCG1, KMCG3, and KMCG6), showing a progressive decrease in consortia complexity (richness) over generation time (Fig. 2a). Ten, nine and five different classes were observed from KMCG1, KMCG3, and KMCG6, respectively. At the genus level, there were 44 and 31 genera classified from KMCG1 and KMCG3, and 14 from KMCG6. Only seven known genera *(Pseudochrobactrum, Chryseobacterium, Lysinibacillus, Acinetobacter, Buttiauxella, Stenotrophomonas, and Comamonas)* and one unclassified were detected with a relative abundance > 0.1 *%* from KMCG6 (Fig. 2b). A similar trend was observed at the most sensitive OTU level, as the observed richness decreased from 162 (KMCG1) to 85 (KMCG6), implicating that several OTUs were affiliated per genera. Seven OTUs were classified to *Pseudochrobactrum,* which was the most diverse genera, followed by *Chryseobacterium and Stenotrophomonas,* both containing four OTUs. Notably, 61.61% of the total sequences were clustered to OTU_457, representing the *Chryseobacterium sp. KMC2 (CKMC2)* in the KMCG6. *Pseudochrobactrum sp. KMC2 (PKCM2)* and *Acinetobacter sp. KMC2 (AKMC2)* accounted for 11.28 % and 5.04 %, respectively. These results show that selection reduced diversity and specific strains were enriched in the consortia.

**Fig. 2.**
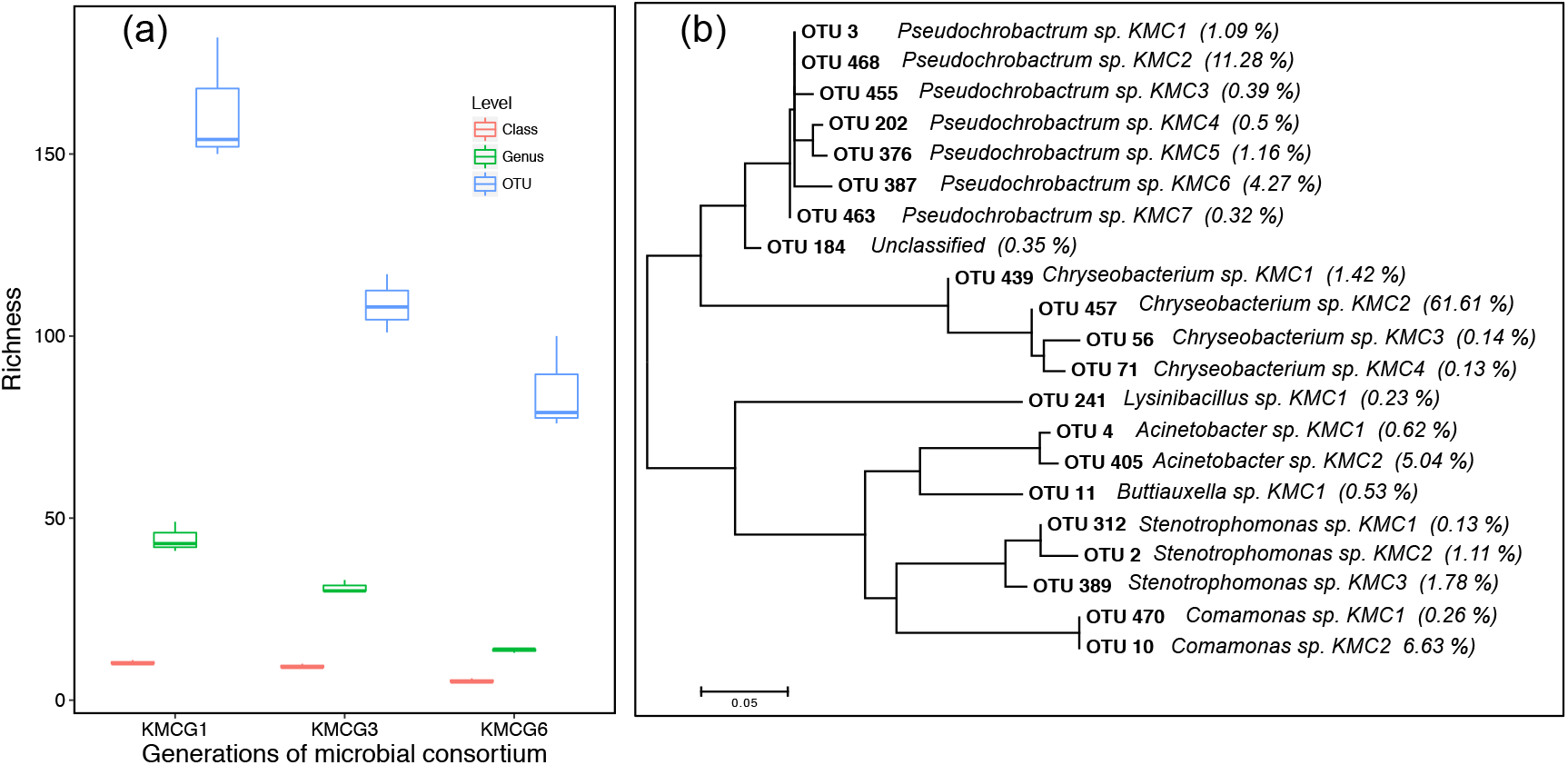
Microbial diversity from the enrichment process. (a). The observed values of class, genus, and OTU from different generations in the enrichment process (KMCG1, KMCG3, and KMCG6). (b). Phylogenetic tree of dominant OTUs from KMCG6 (relative abundance >0.1 %).

Note that it is preferable to obtain microbial consortia with desired functions prior to practical implementation, which can be obtained from naturally occurring microbial communities. But in most cases, this is still not sufficient to reach a desired effective consortium with a low diversity, which can be further used for downstream industrial applications (28, 29). For instance, more than 85 potential strains were detected from KMCG6, although the actual amount may be lower because multi-OTUs could originate from a same single strain due to intrinsic variability amongst multiple copies of 16S rRNA gene (30, 31). KMCG6 still had a high diversity with 14 genera and more than 21 dominating OTUs (> 0.1 %) with distinct relative abundance, which likely corresponded to strains with different functionality towards keratin degradation. It indicates that the community diversity was still too high for understanding the underlying mechanisms. Therefore, simplification of this system is required to obtain more controllable consortia for downstream keratinous waste management, and with potentially enhanced activity.

### Optimal dilution for construction of a library of SMC

The dilution-to-extinction strategy has already proven its efficiency for obtaining functional isolates and microbial consortia from various initial environmental inoculums such as seawater and rumen liquor (32–34), which was adapted here to our pre-enriched KMCG6. In order to determine the optimal dilution to KMCG6, we performed gradient dilutions with 24 replicates in keratin medium (KM). Cell density, protease, and keratinase activities measuring from individuals were used to evaluate the dissimilarity between distinct dilutions. Approximately 21 % (5) and 92 % (22) of the diluted replicates displayed no cell growth from dilution 10^−9^ and 10^−10^, respectively. Dilution 10^−10^ was excluded based on the observed low growth and lack of sufficient degradation efficiency. Comparison of different dilutions was calculated according to their characteristics of degrading capacities (cell growth and enzyme activities) with Euclidean distance (Fig. 3a). It suggests that the profiles of dilution 10^−2^ to 10^−8^ had high similarity. Notably, dilution 10^−9^ shows a clear and expected higher variability compared to lower dilutions (PERMANOVA, FDR < 0.1), such as e.g. 10^−8^, which has a good potential for assembling effective SMC.

**Fig. 3.**
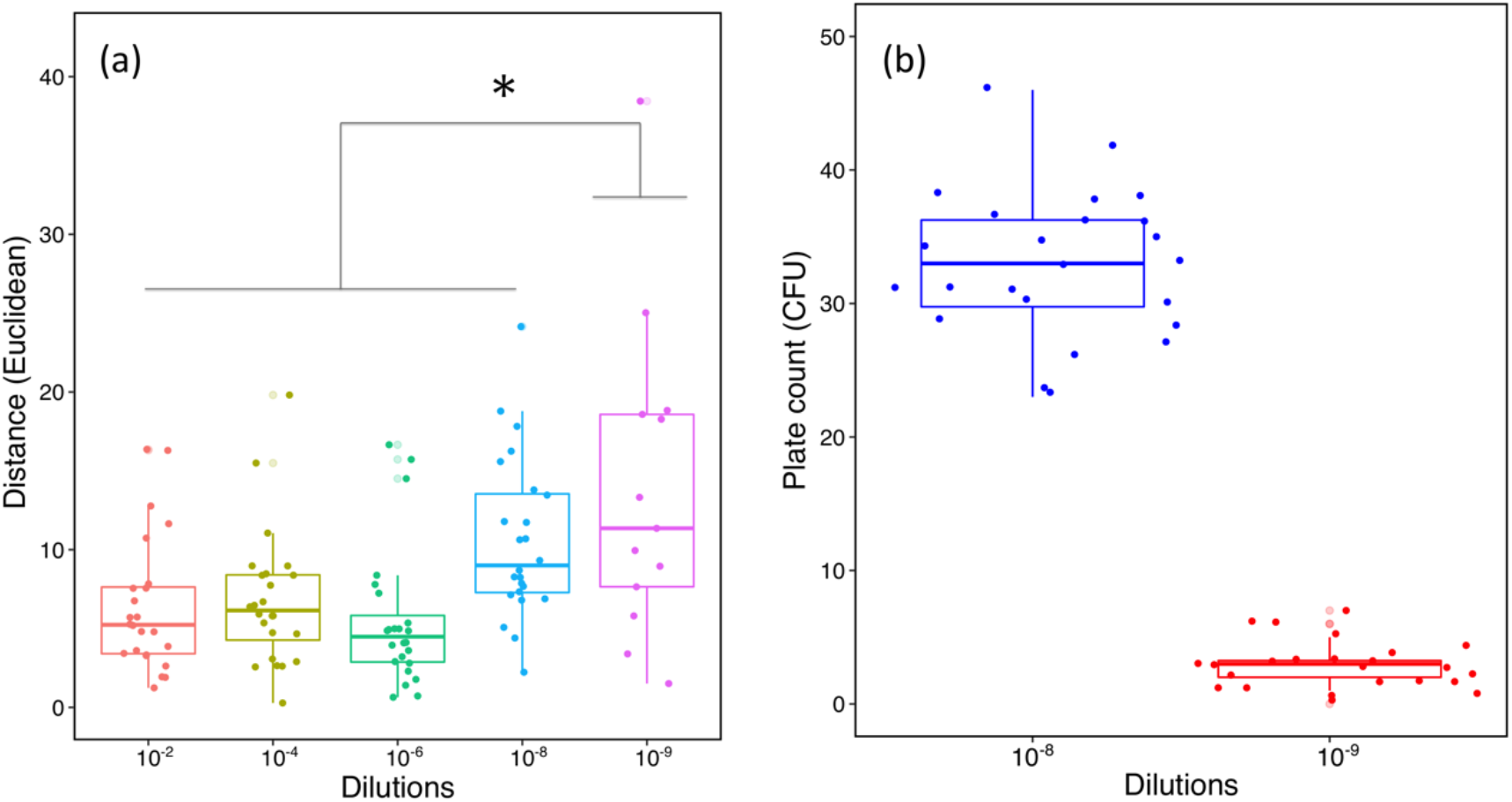
Characteristic comparison of SMC from the dilution-to-extinction culture. (a). Distance comparison based on the characteristics of OD and protease and keratinase activity, with Euclidean dissimilarity index (n=24). Star indicates significant statistical difference (PERMANOVA, FDR < 0.1). (b). Numbers of CFU from dilution 10^−8^ and 10^−9^ by plate counting (n=24).

Once an efficient pre-enriched consortium is secured, determining optimal dilution is as a critical step to obtain good heterogeneity in the subsequent SMC. Here, serial dilutions were carried out to simplify KMCG6. A comprehensive comparison was done for all dilutions, including 24 replicates per dilution. Concerning the number of replicates included, a sufficient quantity provides more reliable and statistically sound results. Previous studies showed that few replicates cultured from distinct dilutions resulted in a limited heterogeneity of functional microbial consortia (33, 35). The higher the better, as it will give greater chance to reproduce and/or even improve the efficiency observed. However, a parsimonious approach would be to rely on an adjusted number of replicates, estimated based on prior evaluation of the taxonomic compositions and abundance of the constituting microbial members in the pre-enriched community.

In addition, we performed CFU counting on LB agar to verify the number of viable cells between dilution 10^−8^ (mean = 33 CFU) and dilution 10^−9^ (mean = 3 CFU), showing an expected decrease of 10-fold (Fig. 3b). More importantly, they possessed various degradative capacities, including high performance in KM. Dilution 10^−9^ hereby was selected as an optimal dilution to further construct a library of SMC consisting of 96 replicates (Fig. S1, for reviewers only). Additionally, 18 SMC were selected for further functional assessment with additional three SMC from dilution 10^−8^.

### Diversity and structure of SMC during keratin degradation

The taxonomic classification of these selected SMC was observed at OTU level using 16S rRNA gene amplicon sequencing (Fig. 4a). In total, 15 OTUs were found in the SMC with a relative abundance above 0.1 %, including 12 observed in KMCG6. Three new detected members were *Chryseobacterium sp. KMC5, Pseudomonas sp. KMC1* and *Stenotrophomonas sp. KMC4.* SMC were clustered into seven groups according to their taxonomic composition (weighted UniFrac). Three SMC from dilution 10^−8^: SMC1, SMC2, and SMC3 were clustered as group 1 (Gr1), all featuring the dominant *CKMC2* (> 62.9%). The relative abundance of *Stenotrophomonas sp. KMC3* and *PKMC2* were more than 5 %. As expected, the composition of SMC in dilution 10^−9^ was set apart from 10^−8^, showing a substantial difference between these two dilutions. 18 SMC were divided into six groups (Gr2 – Gr7), which had heterogeneous profiles in terms of OTU diversity (OUTs = 2-12). 14 SMC (SMC4 – SMC17) from dilution 10^−9^ still featured the dominant *CKMC2,* which initially constituted the majority of KMCG6. Four SMC (SMC18 – SMC21) contained little to none of *CKMC2* and they were classified into two different groups (Gr6 and Gr7).

**Fig. 4.**
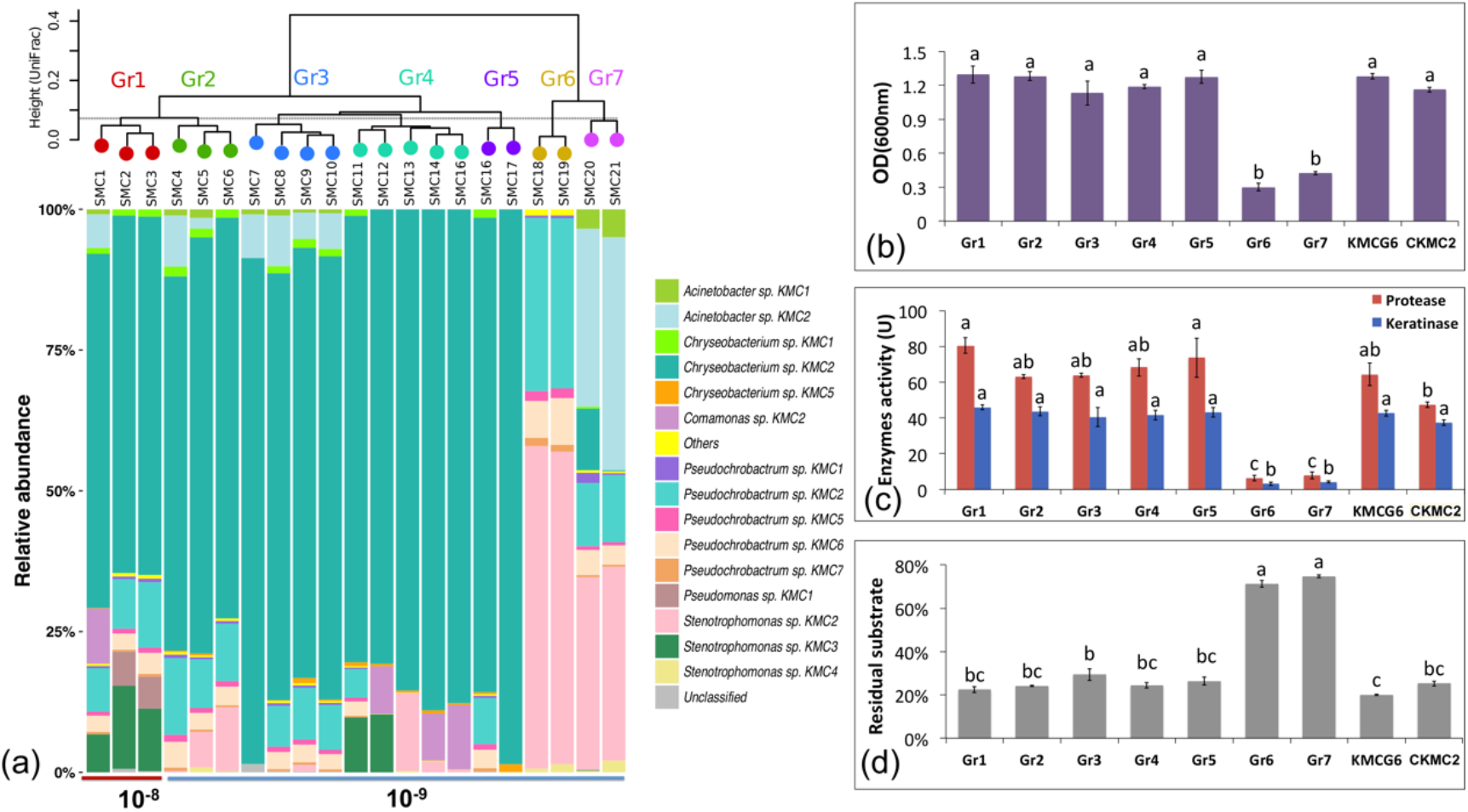
Integrated comparison of selected simplified microbial consortia. (a). 21 SMC divided into seven groups based on their community similarities from 16s rRNA gene analysis (weighted UniFrac distance). Group 1 (Gr1) includes SMC1, SMC2, and SMC3 from dilution 10^−8^. 18 selected SMC (SMC4 – SMC21) were clustered as six groups (Gr2 – Gr7) from dilution 10^−9^. (b). Cell density (OD_600nm_) comparison of group 1-7, KMCG6 and the single strain *Chryseobacterium sp. KMC2 (CKMC2)*. (c) Comparison of enzymes (protease and keratinase) activity. (d) Comparison of the residual substrate ratio. Lowercase letters (e.g. a, b and c) in 4b, 4c and 4d refer to significant differences between groups with one-way ANOVA using post-hoc Tukey’s HSD test (*p* < 0.05).

It is a remarkable fact that several phylogenetically-related strains from the same species coexist in the KMCG6 (Fig. 2b), which is consistent with another similar enrichment study (28). The functions of individual strains are likely to be diverse within species and it is an unresolved challenge to identify the intraspecific variablility (36). Here taxonomic classification of most individual SMC showed that only a few strains belonging to same species remained, which demonstrates that the method conveniently selected a simplified microbial consortium with specific functional strains, avoiding the ones with undesired traits. Besides, strains with low abundance emerged when compared to the initial microbial consortium (KMCG6), suggesting the approach is also efficient to acquire potentially crucial rare species into the SMC.

### Comparative analysis of degradative capacities

The initial microbial community is likely to be divided into different functional groups along with the gradual dilution process due to random reassembly of microbes caused by extinction and sampling effects (23). Cell density was measured at day three for all SMC along with protease and keratinase activities. This timeframe is optimized to achieve the highest cell densities and enzyme activities according to our previous work (11). The keratinolytic characteristics of SMC were measured individually, and displayed on the basis of the defined groups by consortia composition (Fig. 4b, 4c and 4d). Additionally, KMCG6 and the *CKMC2* isolate were compared to these groups. These results clearly show two distinct performance categories present amongst the groups in term of cell density, enzyme activity, and residue ratio. The first category includes mostly groups from dilution 10^−9^: Gr2, Gr3, Gr4, and Gr5, together with Gr1 (10^−8^), all displaying good performance similar to KMCG6, while Gr6 and Gr7 have low performance with weak keratinolytic activity. SMC from 10^−8^ (SMC1, SMC2, and SMC3) show good growth rates, with OD_600nm_ reaching up to (1.15 – 1.3). The OD_600_ of the 18 SMC from dilution 10^−9^ ranges from 0.1 to 1.2. No visible difference in cell growth is observed between the good performers and KMCG6. It is worth noticing that the protease activity from Gr5 was significantly higher than *CKMC2.* On the other hand, the residue ratio of SMC from dilution 10^−9^ was very divergent, indicating that the dilution-to-extinction cultures have succeeded in generating heterogeneous functional SMC.

### Potential biotic interactions

To enhance the association between strains and degradative capacity, a Pearson’s correlation based network was adopted (Fig. 5). As expected, the cell density and enzyme activities had a significant positive correlation with degradation efficiency. Seven strains *(CKMC2, PKMC2, Psedochrobacterum sp. KMC5, Psedochrobacterum sp. KMC6, Psedochrobacterum sp. KMC7, Stenotrophomonas sp. KMC2 (SKMC2) and Stenotrophomonas sp. KMC4*) were connected with keratinolytic activity, which are likely to be the key players in the keratin degradation. Among them, *CKMC2* was the only strain that had positive correlations with all of the keratinolytic characteristics, suggesting that *CKMC2* is the keystone strain in the degradative process. Interestingly, all of the other strains in this correlated network have negative correlations with *CKMC2.* However, *PKMC2* has also a positive correlation with the degradation efficiency while having a negative one with *CKMC2,* indicating that these two strains likely undergo competitive exclusion, leading to the dominance of *CKMC2* (Fig. 4a). *PKMC2* also has numerous positive correlations with other strains such as *SKMC2* and *Pseudochrobactrum sp. KMC7*, indicating that *PKMC2* may play a significant role to maintain the structure of the microbial consortia. However, *SKMC2* is associated to a decrease in degradation efficiency, indicating that it may be a cheater strategist benefiting from *PKMC2.* Additional four strains *(Acinetobacter sp. KMC1, AKMC2, Stenotrophomonas sp. KMC3,* and *Pseudomonas sp. KMC1)* only had satellite correlations, which probably have no effect on keratin degradation and could therefore be removed from the community. The network analysis based on the SMC library can be used to identify actual microbial interaction at the strain level, which may help to further optimize functional microbial consortia.

**Fig. 5.**
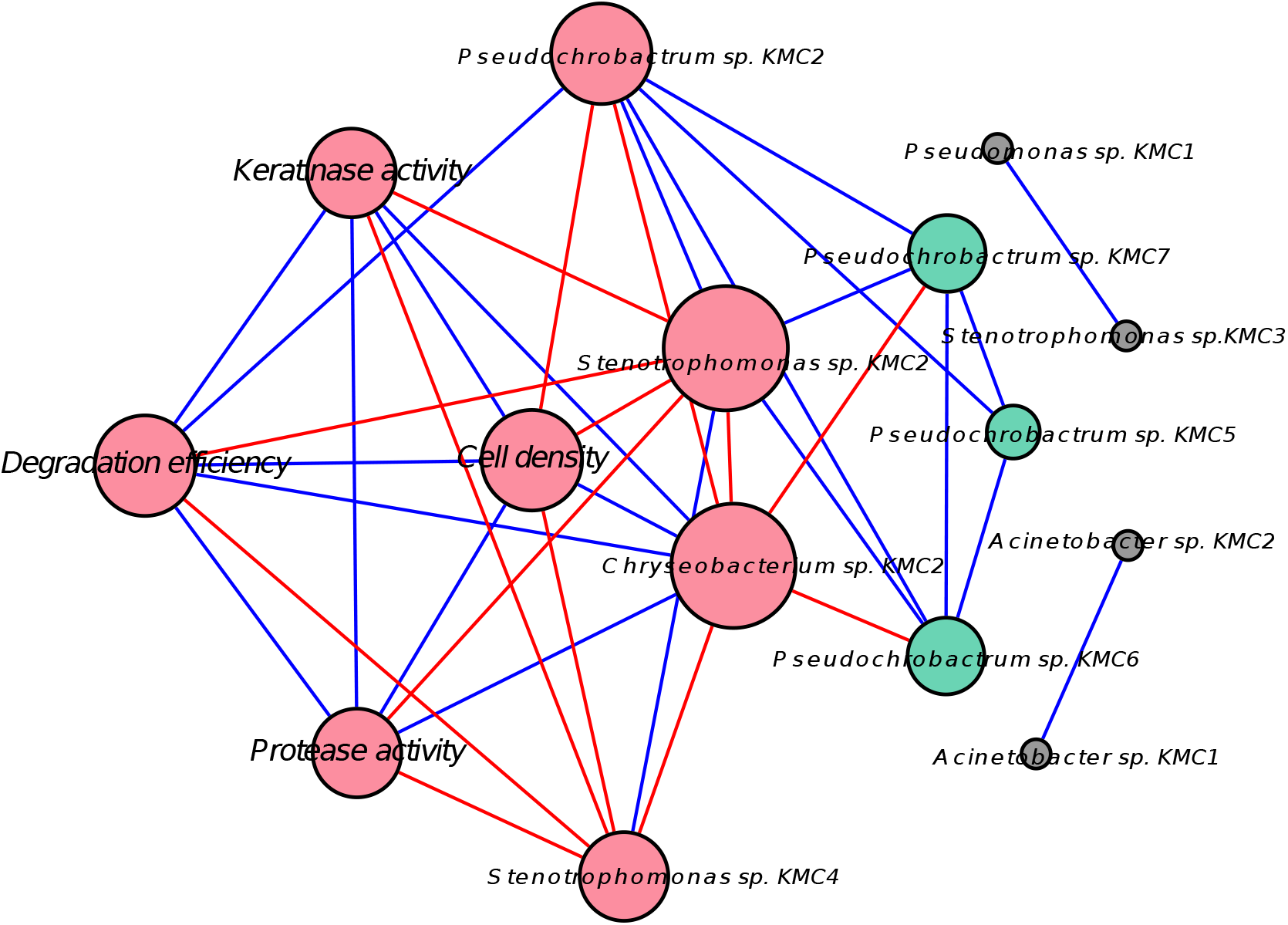
Network showing relationships between strains and degradative capacity using Pearson’s correlation. The nodes represent either OTUs or characteristics of keratin degradation, e.g. degracation efficiency and keratinase activity. Red nodes indicate the direct correlation with degradation efficiency (DE). Green nodes had an indirect correlation with DE through other strains. Grey nodes had no significant correlation with DE. Size of the nodes reflects the number of connections (degree); larger nodes have more significant connections. Blue lines represent positive correlations and red lines represent negative correlation.

The microbial diversity in the so-obtained SMC decreased remarkably while maintaining an equivalent keratinolytic capacity. It supports that dilution-to-extinction represents a simple and practical strategy, simplifying microbial consortia without losing efficiency. Currently, microbial consortia are not only applied on biodegradation. They have been applied extensively in other fields such as biomining, bioremediation, biofilm formation, or biosynthesis (37–39). We show that dilution-to-extinction culture is efficient to simplify microbial consortia, without impairing keratinolytic activity. Overall, a strategy combining enrichment and dilution-to-extinction cultures was applied successfully to assemble several SMC, which had heterogeneous capacities for keratinous waste management. Meanwhile, it provides a view of potential interactions among strains, which can be used for further designing and engineering of microbial consortia. This approach promotes the efficient selection of simplified functional consortia from high diversity environmental habitats and raises the possibility to obtain satisfactory microbial consortia for practical applications.

## Materials and Methods

### Substrate and medium preparation

Mixed α-keratin materials (raw bristles and hooves) were collected from a Danish Crown slaughterhouse (Bragesvej, Denmark), washed with tap water thoroughly, and cut to about 2 mm in diameter mechanically by Daka Sarval (Løsning, Denmark). Keratin medium (KM) was prepared with 1 % keratinous materials with mineral salt medium (0.5 g/L of NH_4_Cl, 0.5 g/L of NaCl, 0.3 g/L of K_2_HPO_4_, 0.4 g/L of KH_2_PO_4_, 0.1 g/L of MgCl_2_ · 6H_2_O) (40), with keratins being the sole carbon source. KM was sterilized by autoclaving (21 min, 120 °C).

### Gradual dilutions of the microbial community

Pre-enriched microbial consortium “KMCG6” from Kang et al. (11) was obtained after serial enrichments over time in successive generation batches, namely Keratin Microbial Consortia Generation batch number 6: “KMCG6”. KMCG6 was inoculated into the LB liquid medium for overnight cultivation with shaking until the optical density (OD_600_) reached 0.7 – 0.8 (250 rpm, 24 °C). The cell suspension was gradually diluted in LB liquid medium with six dilutions (10^−2^, 10^−4^, 10^−6^, 10^−8^, 10^−9^ and 10^−10^). For each dilution 24 replicates were prepared by transferring 200 μL into individual wells of a 96-well plate. Plates were incubated overnight (250 rpm, 24 °C). Subsequently, all replicates from different dilutions were inoculated into 24-well plates with 1.5 mL KM at 1:100 ratio, and grown for three days (250 rpm, 24 °C) before functional assessment *via* cell density and protease acitivity assays.

### Cell density measurement and cell number count. (i) Cell density measurement

Cultures were left standing for 10 min. to precipitate large suspended keratin particles after cultivation. Thereafter, 200 μL cell suspension from all dilutions were transferred to 96-well plates. Cell density was measured (OD_600_) using a microplate reader (Biotek, ELx808). **(ii) Cell number count.** 200 μL cell suspension from 24 replicates of dilutions (10^−8^ and 10^−9^) was plate spread on LB agar plates. Cell numbers were counted according to observable colony-forming units (CFU) on the plates after 48 h.

### Construction of various microbial consortia library

A total of 96 SMC were obtained based on optimal dilution condition (10^−9^) and 18 SMC were further selected from the library after the functional assessments. In addition, three SMC from dilution 10^−8^ were as the control. In order to evaluate the degradation capacities accurately, the cultivation of 21 SMC was scaled up to 100 mL KM by 1:100 ratio for 5 days (200 rpm, 24 °C).

### Enzyme activity assays. (i) Protease activity assay

Protease activity of microbial consortia was assessed with azocasein (Sigma-Aldrich, St. Louis, MO, USA) as described previously (11). 100 μL supernatant with 50 μL 1% (w/v) azocasein were incubated at 30 °C for 30 min. with shaking at 200 rpm, then stopped by adding 150 μL 10% (w/v) trichloroacetic acid and incubated at 4 °C for 15 min. 100 μL mixture was mixed with 100 μL of 0.5 M NaOH after centrifugation. Absorbance was recorded in 96-well plates at 415 nm and one unit (U) of protease activity was defined as 0.01 increase in absorbance. **(ii) Keratinolytic activity assay**. Preparation procedure for azokeratin and associated keratinolytic activity assay were described previously (11). The prepared keratin materials were coupled with a diazotized aryl amine to produce a chromophoric derivative, sulfanilic acid azokeratin. Keratinase activity of microbial consortia was assayed using azokeratin as a substrate. The reaction mixture of supernatant and azokeratin were incubated for 1 h at 30 °C with shaking at 200 rpm, then cooled to room temperature for 5 min. 200 μL supernatant without azokeratin was transferred to 96-well plates and absorbance was measured as above. One unit (U) of keratinase activity was defined as the amount of enzyme required for 0.01 increasing in absorbance.

### Residual keratin substrate weight

Keratin residue was collected from all of the microbial consortia after five days of cultivation in KM. The residual substrate was washed with deionized water using vacuum filtration until the flow-through was colourless to remove all of the microbial biomass and then dried at 50 °C for 48 h. The residual substrate was weighed and reported as the percentage (w/w %) of initial keratin substrate.

### Composition analysis of the microbial consortia

A total of 5 mL cell suspension was collected from each microbial consortium after three days of cultivation. DNA extraction was done using the FAST Soil DNA Kit (MP Biomedicals, U.S.A), following the manufacturer’s instructions. PCR amplification and sequencing preparation were performed as previously described (41), using the primers Uni341F (5’-CCTACGGGRBGCASCAG-3’) and Uni806R (5’-GGACTACHVGGGTWTCTAAT-3’) flanking the V3 and V4 regions of the 16S rRNA gene (42, 43). Purification of PCR products was done with Agencourt AMPure XP Beads (Beckman Coulter Genomics, MA, USA) according to the manufacturer’s instructions. They were further quantified using Quant-iT™ High-Sensitivity DNA Assay Kit (Life Technologies) and pooled in equimolar concentrations using SequalPrep™ Normalization Plate (Thermo Fisher Scientific) before concentration using the DNA Clean and Concentrator™-5 kit (Zymo Research, Irvine, CA, USA). Finally, a 20 pM pooled library was subjected to paired-end (2×250 bp) high-throughput sequencing on an Illumina MiSeq platform (Illumina, San Diego, CA, USA) using MiSeq reagent kit v2. Raw sequencing data were handled as previously described (44), which included quality control, trimming and clustering of sequences into OTU using a 97 % pairwise sequence similarity threshold. A representative sequence from each OTU cluster was given taxonomical annotation using RDP database (45). A read contingency table with OTU information including enrichment process was exported at the species level. The comparison of KCCG1, KMCG3, and KMCG6 was performed at class, genus and OTU levels. Phylogenetic tree of KMCG6 was constructed based on OTU sequences using MEGA7 (46) by maximum likelihood method. Only OTU with a relative abundance above 0.1 % were considered. The raw sequencing data is being prepared for deposition at the NIH short read archive.

### Strain identification

SMC from dilution 10^−9^ with high keratinolytic activity were plated on LB plates and incubated at 24 °C for 48 h. In order to secure strain purity for accurate identification and isolation, single colony was picked and inoculated into LB liquid medium for overnight cultivation until OD_600_ reached 0.7 – 0.8 (250 rpm at 24 °C), then plated on new LB plates. This procedure was repeated three times until all colonies on the LB plates had the same morphological characteristics. Afterward, one single colony was picked, cultured in LB liquid medium overnight. DNA was extracted from 2 mL culture with FAST Soil DNA Kit (MP Biomedicals, U.S.A), following the manufacturer’s instructions. The DNA was used as the template with primers 27F (5’-AGAGTTTGATCMTGGCTCAG-3’) and 1492R (5’-TACGGYTACCTTGTTACGACTT-3’) to amplify the full 16S rRNA gene, followed by BLAST identification with assembled OTUs.

### Statistical analysis

Euclidean distance was calculated to determine the functional dissimilarity of SMC from different dilutions in terms of cell density, enzymes activity, and residual ratio. The multivariate homogeneity of group dispersion was analysed with “betadisper” (R-package “vegan”) and assessed using PERMANOVA. Benjamini–Hochberg correction was applied to control false discovery rate (FDR) for multiple testing (47). The significance level within dilutions was defined as the FDR < 0.1. For the selected 21 SMC, grouping was performed according to the compositional similarity using weighted UniFrac distance metric (48). Statistical differences of degrading capacities among SMC groups, KMCG6, and single strains were performed by one-way ANOVA using post-hoc Tukey’s HSD test (*p* < 0.05). Relationships between microbes and degrading capacities were evaluated by using Pearson’s correlation, and visualized with Gephi (49). All of the statistical analysis in this study was caculated by using R (version 3.5.0).

## Acknowledgments

This research was funded by the Danish Innovation Fund (grant number 1308-00015B, Keratin2Protein) and also under the support of the Chinese Scholarship Council (CSC) Scholarship Program.

